# *Chryseobacterium indologenes* mediates resistance to osimertinib by activating the IGF1R pathway in EGFR-mutant lung adenocarcinoma

**DOI:** 10.1101/2025.10.08.681060

**Authors:** Wendong Li, Keqiang Zhang, Mahima Raul, Aviva Rotter-Maskowitz, Deborah Nejman, Jan Potempa, Ravid Straussman, Dan J. Raz

## Abstract

**Introduction:** Lung cancers harboring sensitizing epidermal growth factor receptor (EGFR) mutations typically exhibit an initial response to osimertinib; however, the development of resistance is inevitable, and there are currently no approved targeted therapies available once resistance emerges. Accumulating evidence indicates that intra-tumoral bacteria can influence tumor biology and contribute to therapy resistance in several cancer types, including pancreatic and colorectal cancer. Despite these findings, whether intra-tumoral bacteria play a role in modulating response and resistance to osimertinib in EGFR-mutant lung adenocarcinoma remains largely unexplored. This study aims to investigate the contribution of intra-tumoral bacteria to osimertinib resistance, thereby providing new insights into resistance mechanisms and identifying potential therapeutic strategies.

**Methods:** Bacteria previously identified in lung cancer tissue samples were cultured in liquid growth medium, and their preconditioned medium (PCM) was collected. PC9 cells were treated with PCM in the presence or absence of osimertinib to screen for bacteria capable of mediating resistance. Cell viability was assessed using Cell Counting Kit-8 assays. To investigate potential mechanisms, Western blotting and receptor tyrosine kinase (RTK) phosphorylation arrays were performed. Control groups included cells treated with osimertinib alone, PCM alone, or vehicle. Statistical analyses were conducted using Student’s t-test as appropriate, with p < 0.05 considered statistically significant. Data represent mean ± SD from at least three independent experiments.

**Results:** The addition of PCM from *Chryseobacterium indologenes* restored cell viability in EGFR-mutant lung adenocarcinoma cells treated with osimertinib. PCM exposure markedly increased insulin-like growth factor-1 receptor (IGF1R) phosphorylation levels. PCM did not enhance cell viability when IGF1R was silenced or inhibited with linsitinib, demonstrating the pathway’s essential role. Proteinase K treatment abolished the ability of PCM to protect cells from osimertinib, and removal of proteins potentially interacting with IGF1R further diminished its efficacy. Finally, PCM also conferred resistance to osimertinib in patient-derived EGFR-mutant lung adenocarcinoma cell lines.

**Conclusion:** Our study demonstrates that the intra-tumoral bacteria *C. indologenes* may significantly influence sensitivity to osimertinib or impart resistance in EGFR-mutant lung adenocarcinoma by activating the IGF1R signaling pathway.

## Introduction

Somatic mutations in the tyrosine kinase (TK) domain of the epidermal growth factor receptor (EGFR) represent the most common actionable alterations in non-small cell lung cancer (NSCLC), occurring in ∼15% of Caucasian and up to 50% of Asian patients with advanced disease.^1, 2^ These mutations are predictive biomarkers for response to EGFR tyrosine kinase inhibitors (TKIs), which are a cornerstone of targeted therapy in NSCLC. Approximately 90% of EGFR mutations are either exon 19 deletions or the L858R substitution in exon 21.^2^ Osimertinib, a third-generation EGFR-TKI, is the current standard first-line treatment for advanced EGFR-mutant NSCLC, offering a median progression-free survival of 19 months.^3, 4^ Despite these advances, resistance to osimertinib is inevitable, and no approved targeted therapies exist once resistance develops.

Multiple mechanisms of osimertinib resistance have been described^5-11^, though they remain incompletely understood. EGFR-dependent mechanisms include loss of the T790M mutation, emergence of tertiary mutations such as C797S, and EGFR amplification.^12^ EGFR-independent mechanisms are more commonly observed after first-line osimertinib therapy and include bypass signaling through pathways such as MET amplification, HER2 amplification, PI3K activation, or RAS-MAPK signaling.^6, 7, 13-23^ Other mechanisms involve downstream alterations such as cyclin D amplification, oncogenic fusions, and histological transformation to small-cell lung cancer.^7 24^ Collectively, these mechanisms highlight the complexity of osimertinib resistance and the urgent need for novel therapeutic strategies.

Beyond genetic and signaling alterations, the tumor microbiome has emerged as an additional factor influencing cancer therapy. Intra-tumoral bacteria have been detected in multiple malignancies, including breast, lung, pancreatic, ovarian, and melanoma tumors^25-31^, with each cancer type exhibiting a distinct microbial composition.^32^ Recent studies demonstrate that tumor-associated bacteria can mediate drug resistance. For instance, γ-proteobacteria in pancreatic tumors metabolize gemcitabine through bacterial cytidine deaminase, inactivating the drug. In lung cancer, Nejman et al. identified diverse bacterial species across more than 400 resected specimens, with variation by histological subtype and smoking status.^33^ However, whether intra-tumoral bacteria contribute to resistance against osimertinib in EGFR-mutant lung cancer remains unknown.

In this study, we identified *Chryseobacterium indologenes*, a gram-negative bacterium previously reported in lung tumors^32^, as a mediator of resistance to EGFR-TKIs, including osimertinib. Although *C. indologenes* is typically non-pathogenic, it can cause opportunistic infections in immunocompromised individuals^34^, partly due to secreted proteases.^35^ Here, we demonstrate that *C. indologenes* promotes osimertinib resistance in EGFR-mutant lung adenocarcinoma cell lines and patient-derived samples via activation of the insulin-like growth factor-1 (IGF1R) signaling pathway, revealing a novel microbiome-driven mechanism of therapeutic resistance.

## Materials and Methods

### Preparation of bacterial pre-conditioned medium (PCM)

Direct long-term co-culture of bacteria with cancer cells was not feasible due to bacterial overgrowth; therefore, bacterial culture broth was used as PCM to test its effect on drug susceptibility. Bacteria were grown overnight at 30°C in nutrient broth (8 g/L in distilled water) until reaching an optical density (OD) of 1. Cultures were then diluted 1:100 and allowed to grow to their logarithmic-growth state until the OD reached 0.3. The bacterial suspensions were then centrifuged at 4,000x g for 20 min, and the supernatants were collected and filtered through 0.22µm filters to obtain sterile PCM, which was used immediately or stored at -80°C until use. Sterile bacterial medium was used as a background (negative) control and subjected to the same processing as the cultures.

### Bacterial culture and rescue screen

A wide phylogenetic panel of 50 bacterial species was grown overnight, each species in its own optimal growth media. Bacterial PCM was obtained for each species as described above. EGFR-mutant lung adenocarcinoma cell lines (LACs; PC9 and HCC4006) stably expressing enhanced green fluorescent protein (eGFP) were used for screening. On day 0, 2,000 LAC cells per well in 40µl were plated in 384-well plate clear-bottom plates (Corning Inc., Corning, New York, USA) using an EL406 liquid dispenser (Agilent Technologies, Inc., Santa Clara, California, USA). Throughout the screen, cells were incubated at 37°C in a humidified incubator with 5% carbon dioxide. On day 1, cells were treated with 10µl of freshly prepared bacterial PCM or sterile bacterial medium negative control using the CyBio Well Vario 96/250 Simultaneous Pipettor (Analytik Jena AG, Jena, Germany). After 45 min, cells were treated with 10µl of 6x prepared drug or DMSO control. EGFR inhibitors (osimertinib, neratinib) were tested in this screen. Each treatment condition was performed in quadruplicate. On day 4, the LAC cells were supplemented with fresh medium, and fresh bacterial PCM and fresh drugs were administered. GFP fluorescence was measured on days 1, 4, 6, and 7 using Cytation3 (BioTek). To assess bacterial-mediated chemoresistance, a rescue score was calculated as (*F*_*PO*_−*F*_*O*_)/(*F*_*C*_−*F*_*O*_), where *F*_*c*_ = control fluorescence intensity (DMSO and bacteria culture medium treatment); *F*_*O*_ = fluorescence intensity for osimertinib alone (inhibition condition); *F*_*PO*_ = fluorescence intensity for PCM + osimeritnib (rescue by PCM). The rescue score takes into account the effect of the bacterial PCM on LAC response to the administered drugs. This score penalizes for insufficient drug-mediated killing.

### Validation of C. indologenes’ impact on LAC resistance to osimertinib

To validate the ability of *C. indologenes* to confer resistance to osimertinib in EGFR-mutant LACs, eGFP-tagged PC9 (PC9-GFP) cells were seeded at a density of 5,000 cells per well in 96-well plates. Cells were treated with osimertinib (10nM; Selleck Chemicals, Houston, Texas, USA) in the presence or absence of bacterial PCM. Cell viability was assessed after 3 days using the Cell Counting Kit-8 (CCK-8) assay, and GFP intensity was imaged after 5 days using a fluorescence microscope. Control groups included untreated cells, and cells treated either with PCM alone or osimertinib alone, or both PCM and osimertinib.

### Preparation of PC9-conditioned medium

To generate PC9-conditioned medium, PC9 cells were seeded in 10cm dishes at 2 × 10^6^ cells/dish in RPMI-1640 with 10% FBS. After 24h, the medium was replaced with 10ml fresh medium. Conditioned medium (con-RPMI) was collected 24h later, filtered (0.22µm), and either used immediately or stored at -80°C. To assess whether con-RPMI promotes the growth of *C. indologenes*, bacteria were cultured in RPMI or con-RPMI, and OD values were measured at 0, 3, 5, 7, and 9h.

### LAC cell culture

LAC cells harboring a sensitizing EGFR mutation, including PC9 [EGFR exon 19 deletion (delE746-A750)], HCC827 [EGFR exon 19 deletion (delE746-A750)], HCC4006[EGFR exon 19 deletion (delL747 - E749, A750P)], H1975[L858R+T790M], and H1650 [EGFR exon 19 deletion (del E746-A750)], were cultured in RPMI1640 medium supplemented with 10% FBS and 1% penicillin-streptomycin. Medium was changed every 2-3 days.

### Fresh patient-derived lung cancer cell lines

Fresh patient-derived lung cancer cell lines were established from surgically resected EGFR-mutant lung adenocarcinoma as previously described.^36^ Primary cultures were maintained in DMEM/F12 (Thermo Fisher Scientific, Waltham, Massachusetts, USA) supplemented with 2% B-27 (Thermo Fisher Scientific), 25 ng/mL fibroblast growth factor (FGF; Thermo Fisher Scientific), 25 ng/mL epidermal growth factor (EGF; Thermo Fisher Scientific), 2μg/mL heparin (EDQM, France), and 100 U/mL penicillin/streptomycin (Thermo Fisher Scientific). Two EGFR L858R-mutant patient-derived lung cancer cell lines, COH4 and COH561912^(38)^, were used. For resistance assays, cells were sub-cultured in RPMI1640 with 10% FBS and 100 U/mL penicillin/streptomycin and exposed to PCM with or without osimertinib. The establishment of patient-derived cell lines was approved by the City of Hope National Medical Center Institutional Review Board (IRB#17196).

### Compound treatment and cell viability analysis

LACs or patient-derived lung cancer cells were seeded onto 96-well plate at a density of 5,000 to 10,000 cells/well. Following a 24-hour period, cells were exposed to the drugs osimertinib, erlotinib, neratinib, as well as the IGF1R inhibitor linsitinib and the ERK inhibitor SCH772948, at the indicated concentrations, in the presence or absence of PCM (100X dilution). All PCM used in the subsequent studies was diluted 100-fold with culture medium unless otherwise noted. Cells exposed only to the background control bacterial culture medium were used as controls in this study, unless noted otherwise. A CCK-8 assay kit (Dojindo Laboratories Co., Ltd., Kumamoto, Japan) was used to determine cell viability following the manufacturer’s protocol. Osimertinib, erlotinib, neratinib, linsitinib and SCH772948 were obtained from Selleck Chemicals (Houston, Texas, USA).

### Proteinase K treatment on bacterial PCM

To evaluate whether protein components in the bacterial PCM mediate drug resistance, PCM and background controls were treated with proteinase K (200 μg/ml) at 56°C for 30 min. Proteinase K activity was then inactivated by treatment with phenylmethylsulfonyl fluoride (PMSF, 1 mM) at room temperature for 30 min. As a control, PCM and background controls treated only with PMSF were included to assess possible PMSF effects. Proteinase K–treated PCM was administered to LAC cells in the presence or absence of osimertinib for cell viability assays. Basal medium treated with PMSF served as an additional control.

### Phospho-receptor tyrosine kinase (phospho-RTK) array

To evaluate changes in RTK phosphorylation in LAC cells and LCSCs after treatment with PCM and osimertinib, a Proteome Profiler Human Phospho-RTK Array Kit (ARY001B; R&D Systems, Minneapolis, Minnesota, USA) was used according to the manufacturer’s instructions. After 24h of treatment, cells were harvested and lysed, and 150-300µg of total protein was applied per array. The relative phosphorylation levels of 49 RTKs were detected simultaneously.

### IGF1 and IGF2 protein treatment

To assess whether IGF1R activation mediates resistance in LACs, cells were treated with recombinant human IGF1 (R&D Systems) or IGF2 (R&D Systems) at 0-100 ng/mL for 3 days in the presence or absence of osimertinib. Cell viability was measured using the CCK-8 assay.

### PCM fraction and heat inactivation of PCM

To understand what components in PCM mediate resistance to osimertinib in EGFR-mutant LACs, PCM was separated using a 10kDa ultrafiltration centrifugal filter at 4,000x g for 30 min. Both the filtrate and the retentate were collected for cell viability assays. Unfiltered PCM was tested in parallel as a control. To assess whether heat inactivation influences PCM’s ability to restore cell viability in the presence of osimertinib, both PCM and the bacterial growth medium were heated at 95°C for 30 min before being administered to PC9 cells with osimertinib.

### Western blot analysis

Equal amounts of protein were separated by SDS-PAGE and transferred to polyvinylidene fluoride (PVDF) membranes. After transfer, membranes were incubated overnight with various primary antibodies at 4°C, followed by incubation with secondary antibodies. Primary antibodies against EGFR, pEGFR, mTOR, pmTOR, AKT, pAKT, ERK1/2, pERK1/2, MET, pMET, IGF1R, pIGF1R, and actin were obtained from Cell Signaling Technology, Inc. (Danvers, Massachusetts, USA).

### siRNA-mediated gene knockdown and drug treatment

To determine whether IGF1R or MET activation mediates osimertinib resistance in EGFR-mutant LACs, cells were reverse-transfected in 96-well plates with 5 pmol/well of siRNA targeting IGF1R (SC-29358), MET (SC-29397), or a non-targeting control (SC-37007) using Lipofectamine RNAiMAX (Invitrogen, Waltham, MA). Twenty-four hours post-transfection, the medium was replaced with varying concentrations of osimertinib in the presence or absence of PCM. Cell viability was assessed 3 days later using the CCK-8 assay.

### Removal of IGF1R-binding proteins from PCM

To deplete IGF1R-binding proteins from PCM, His-tagged Dynabeads™ (Invitrogen, Waltham, MA) were incubated with recombinant His-tagged IGF1R protein (ACRO Biosystems, Beijing, China; Cat# IGR-H5229) for 10 min at room temperature to allow binding. PCM was then applied to the IGF1R-bound beads and incubated for 30 min at room temperature. As a control, PCM was incubated with beads lacking IGF1R. Beads were magnetically separated, and the supernatant, now depleted of IGF1R-interacting proteins, was collected and designated as incomplete-PCM.

### Statistical analysis

All experiments were carried out in triplicate. All analyses were performed using GraphPad Prism software, version 9.4.1 (Dotmatics, Boston, Massachusetts, USA). The *t* test was used to determine the statistical significance of group comparisons between LACs treated with osimertinib in the presence or absence of PCM. A *p* value less than 0.05 was considered statistically significant.

## Results

### C. indologenes can confer resistance to osimertinib in EGFR-mutant LACs

We observed a significant elevation in GFP fluorescence intensity in PC9 cells co-treated with PCM and osimertinib compared to those treated with osimertinib alone (Fig. 1A). To validate this observation, cell viability assays using CCK-8 were performed on PC9 cells exposed to the combined regimen of osimertinib and PCM. Osimertinib monotherapy resulted in approximately 50% viability, whereas the addition of PCM restored viability to levels comparable to untreated controls (background control and DMSO without PCM) (Fig. 1B). To demonstrate the dose-dependent effects of PCM in conjunction with osimertinib, a dilution series of PCM ranging from 6-fold to 1,536-fold was administered to PC9 cells alongside osimertinib. CCK-8 results indicated a significant enhancement of cell viability at PCM dilutions up to 800-fold, with decreased viability observed at minimal dilutions (6- and 12-fold), likely attributable to elevated ionic strength of the medium (Fig. 1B). Parallel assays using erlotinib and neratinib yielded comparable results (Fig. 1C, D). Furthermore, PCM at a 100-fold dilution maintained its capacity to increase viability even with escalating osimertinib concentrations up to 400 nM (Fig. 1E). These findings imply that *C. indologenes*-derived PCM reduces PC9 cell sensitivity to osimertinib, erlotinib, and neratinib.

**Figure 1.**
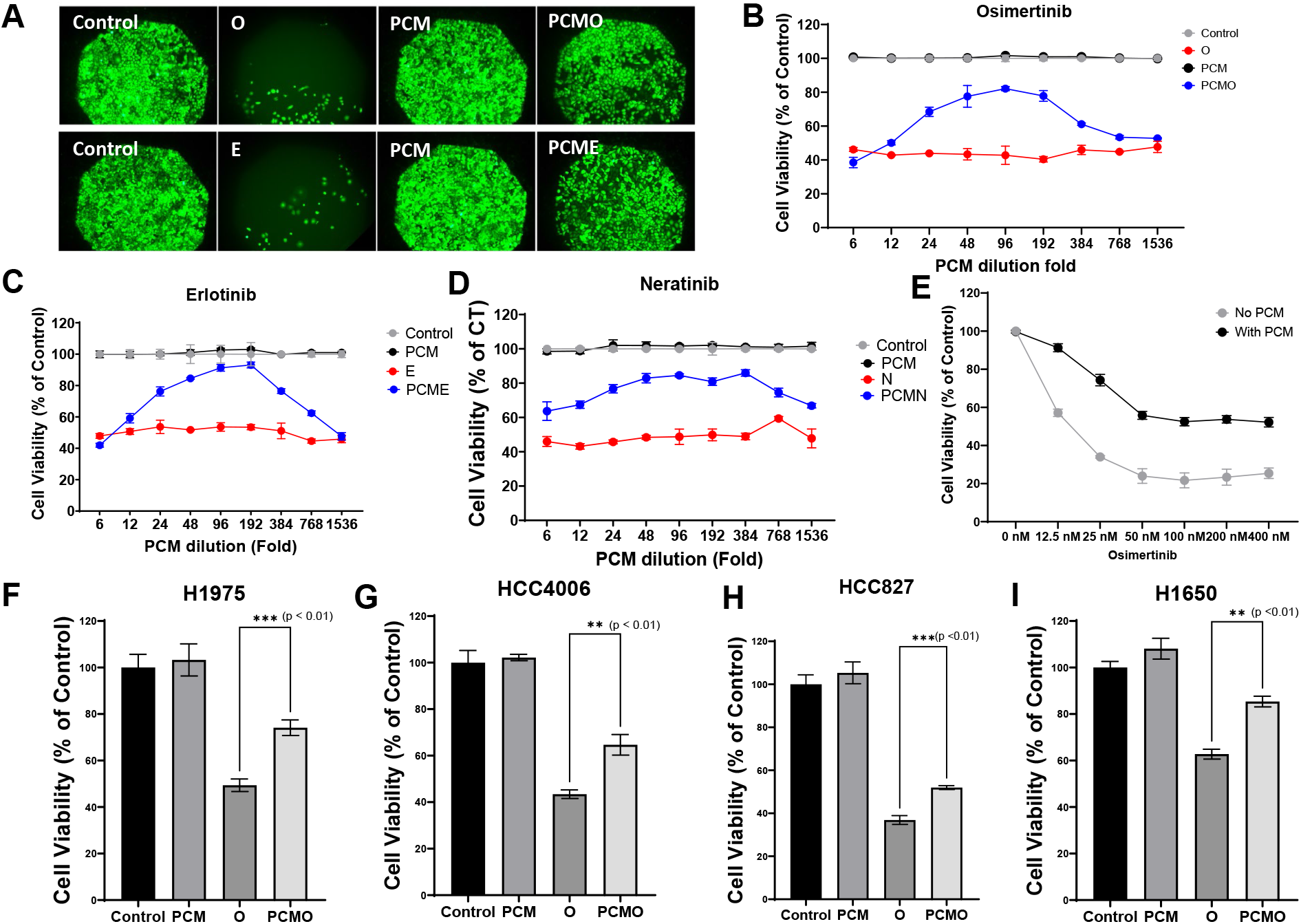
The effect of bacterial PCM (filtered pre-conditioned medium from bacterial culture) on the response of EGFR-mutant lung adenocarcinoma cells to EGFR-TKIs. (A) GFP intensity was scanned after 7 days of treatment. The dose effect of PCM on the response of (B) osimertinib, (C) erlotinib, and (D) neratinib. (E) Effect of PCM on the dose-response of osimertinib on PC9 cell viability. Cell viability assay for (F) H1975, (G) HCC4006, (H) HCC827, and (I) H1650 was performed after 3 days of treatment with osimertinib and PCM.

Extending these analyses to other EGFR-mutant LAC cells, namely, HCC827, HCC4006, NCI-H1650, and NCI-H1975, revealed that PCM co-treatment consistently elevated cell viability in the presence of osimertinib (Fig. 1F-I). Collectively, our results suggest that *C. indologenes* PCM confers resistance to EGFR-TKIs, particularly osimertinib, across multiple EGFR-mutant LAC models.

### Other species of the genus Chryseobacterium can also confer resistance to osimertinib to EGFR-mutant LACs

Given that the genus Chryseobacterium encompasses 112 species^37^, it is plausible that multiple species within this genus possess the capacity to induce resistance to osimertinib in EGFR-mutant LACs cells, similar to *C. indologenes*. To investigate this hypothesis, five commercially available Chryseobacterium species—*C. shigense, C. scophthalmum*, C. *indoltheticum, C. angstadti*, and *C. diehli*—were assessed for their impact on osimertinib sensitivity in PC9 cells. When combined with osimertinib treatment, CCK-8 cell viability assays revealed that *C. indoltheticum, C. scophthalmum*, and *C. shigense* significantly enhanced cell viability, whereas *C. angstadti* and *C. diehli* did not elicit this effect (Fig. 2). These results indicate that resistance to osimertinib in EGFR-mutant LACs is not restricted to *C. indologenes* but can also be mediated by other select species within the Chryseobacterium genus, underscoring the potential broader clinical significance of bacterial-mediated drug resistance mechanisms in this context.

**Figure 2.**
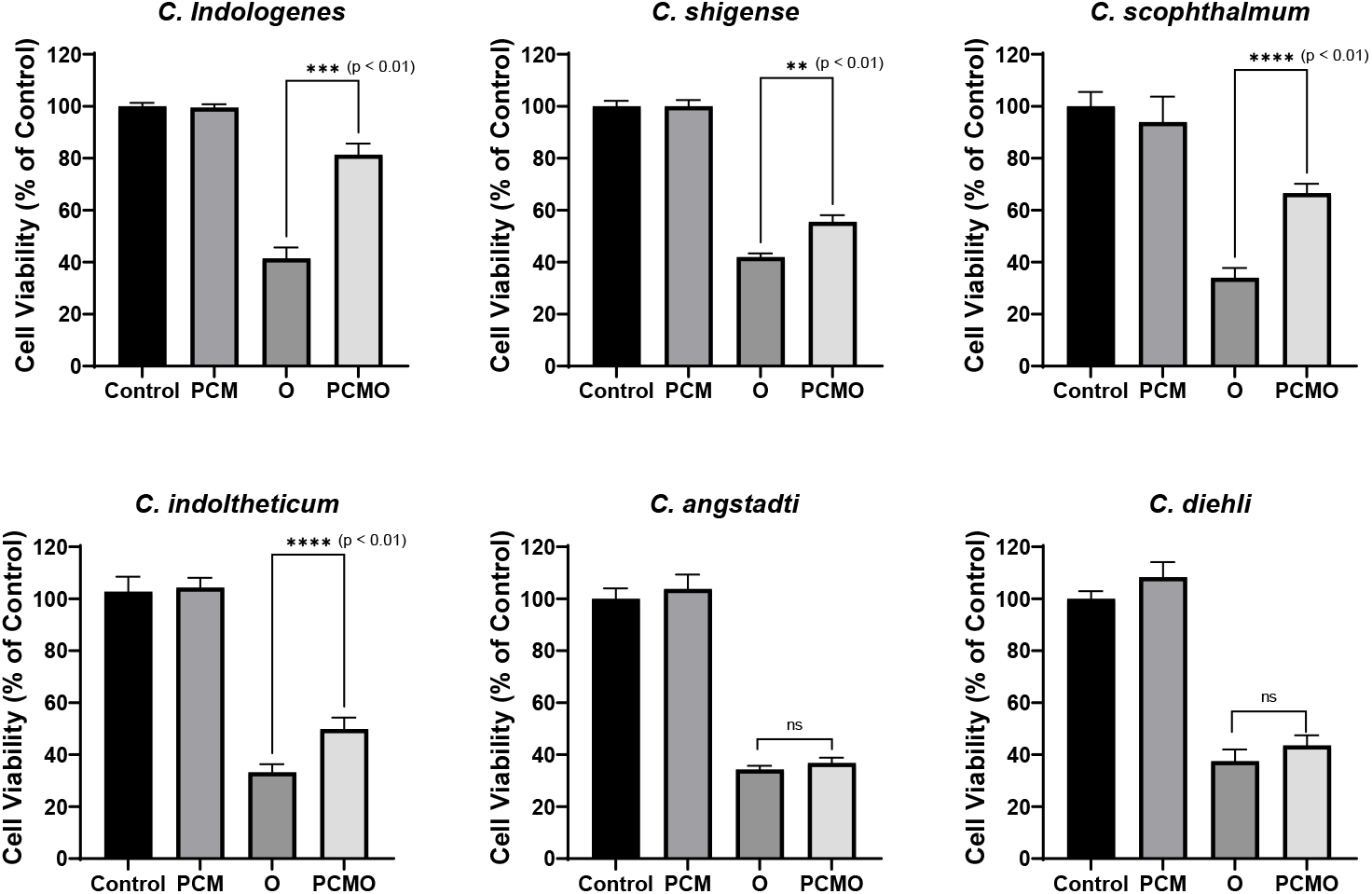
The effect of bacterial PCM from other *Chryseobacterium* strains on the response of EGFR-mutant lung adenocarcinoma cells to osimertinib. Cell viability assay was performed after 3 days of treatment with PCM and osimertinb (10nM) in PC9 cells.

### C. indologenes confers apoptotic resistance to osimertinib and promotes proliferation of EGFR-mutant LACs

To elucidate the effects of *C. indologenes* on osimertinib-induced apoptosis, we conducted flow cytometric analysis of apoptosis in PC9 cells. The addition of *C. indologenes* PCM to osimertinib-treated cells resulted in a marked reduction in apoptotic cell population, indicating that PCM suppresses apoptosis triggered by osimertinib (Fig. 3A). It should be noted that bacterial basal medium without conditioning served as the control. Subsequently, we investigated whether *C. indologenes* infection enhances proliferation in EGFR-mutant LAC cells. Although Fig. 1 reports cell viability rather than direct proliferation, increases in viability upon PCM treatment of PC9, HCC827, and H1650 cells suggest a potential proliferative effect. This was further corroborated by EdU immunofluorescence staining, which demonstrated significantly elevated DNA synthesis in LAC cells exposed to PCM during osimertinib treatment (Fig. 3B, C). Mechanistically, western blot analyses revealed that PCM restores phosphorylation levels of AKT, mTOR, and ERK1/2 proteins, signal transducers known to promote cell proliferation and survival (Fig. 4B). Notably, treatment with the ERK inhibitor SCH772948 (at 100-fold PCM dilution) abrogated PCM-mediated resistance to osimertinib, indicating that ERK pathway activation is critical for this resistance phenotype (Fig. 3D).

**Figure 3.**
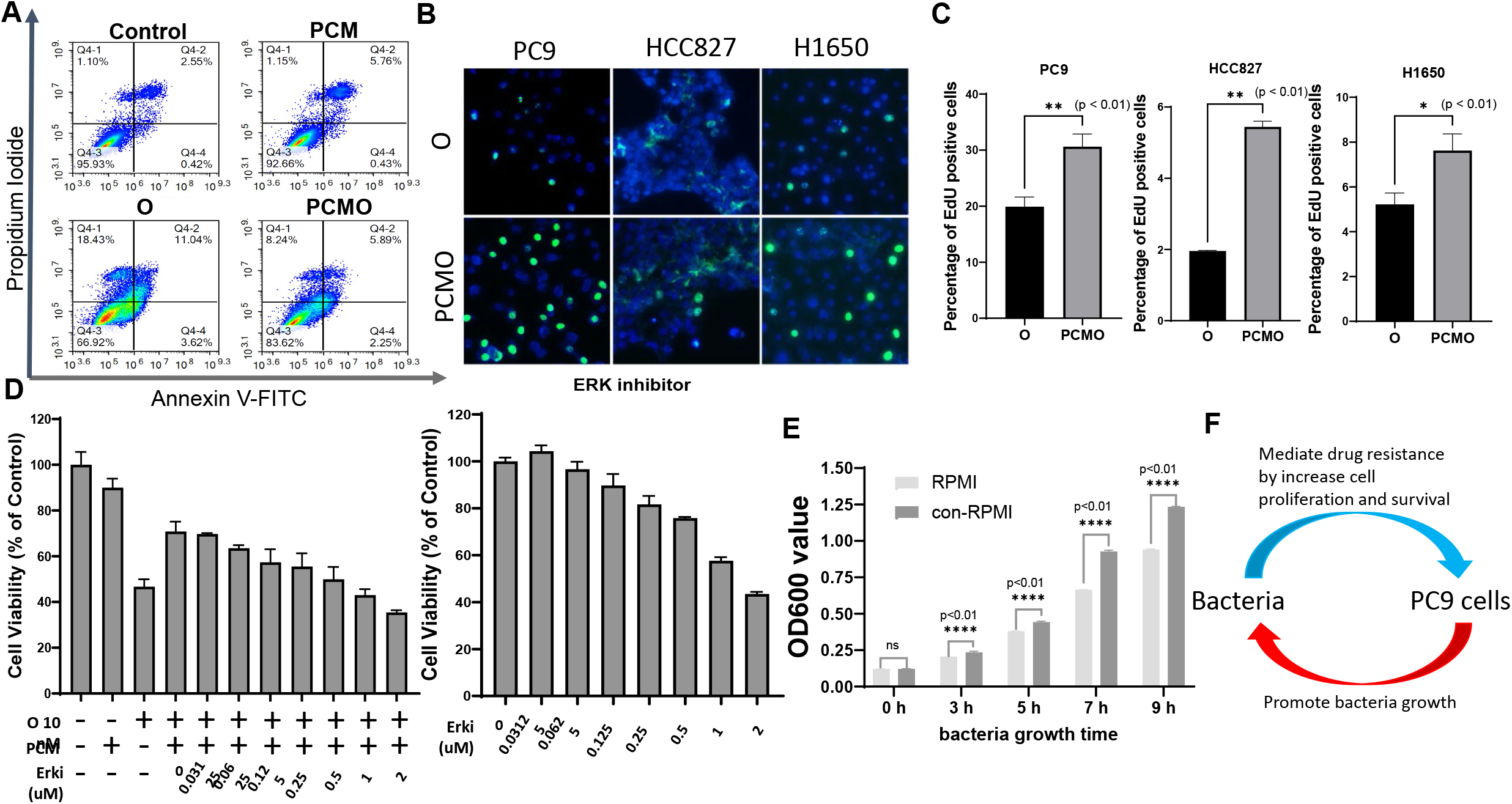
PCM mediates apoptosis resistance and promotes LAC proliferation. (A) Flow cytometry analysis for PC9 cells treated with Osimertinib (20nM) and PCM for 24h. Numbers (%) indicate the percentage of cells stained with Annexin V–FITC- and/or propidium iodide. (B) LAC cells were treated with osimertinib in the presence or absence of PCM for 24h, and then EdU staining was performed. Green denotes EdU-positive cells and blue denotes nuclei. (C) percentage of EdU-positive cells was compared between osimertinib and PCMO. (D) ERK inhibitor SCH772948 abolished PCM-mediated resistance to osimertinib. PC9 cells were treated with the combination of osimertinib, ERK inhibitor and PCM for 72h, and then CCK-8 assay was performed. (E) *C. indologenes* were cultured in PC9-conditioned medium and growth was monitored. Con-RPMI = PC9-conditioned medium, ½ con-RPMI = adding half of amount of con-RPMI. (F) Schematic of the bidirectional interaction between PC9 cells and *C. indologenes*.

**Figure 4.**
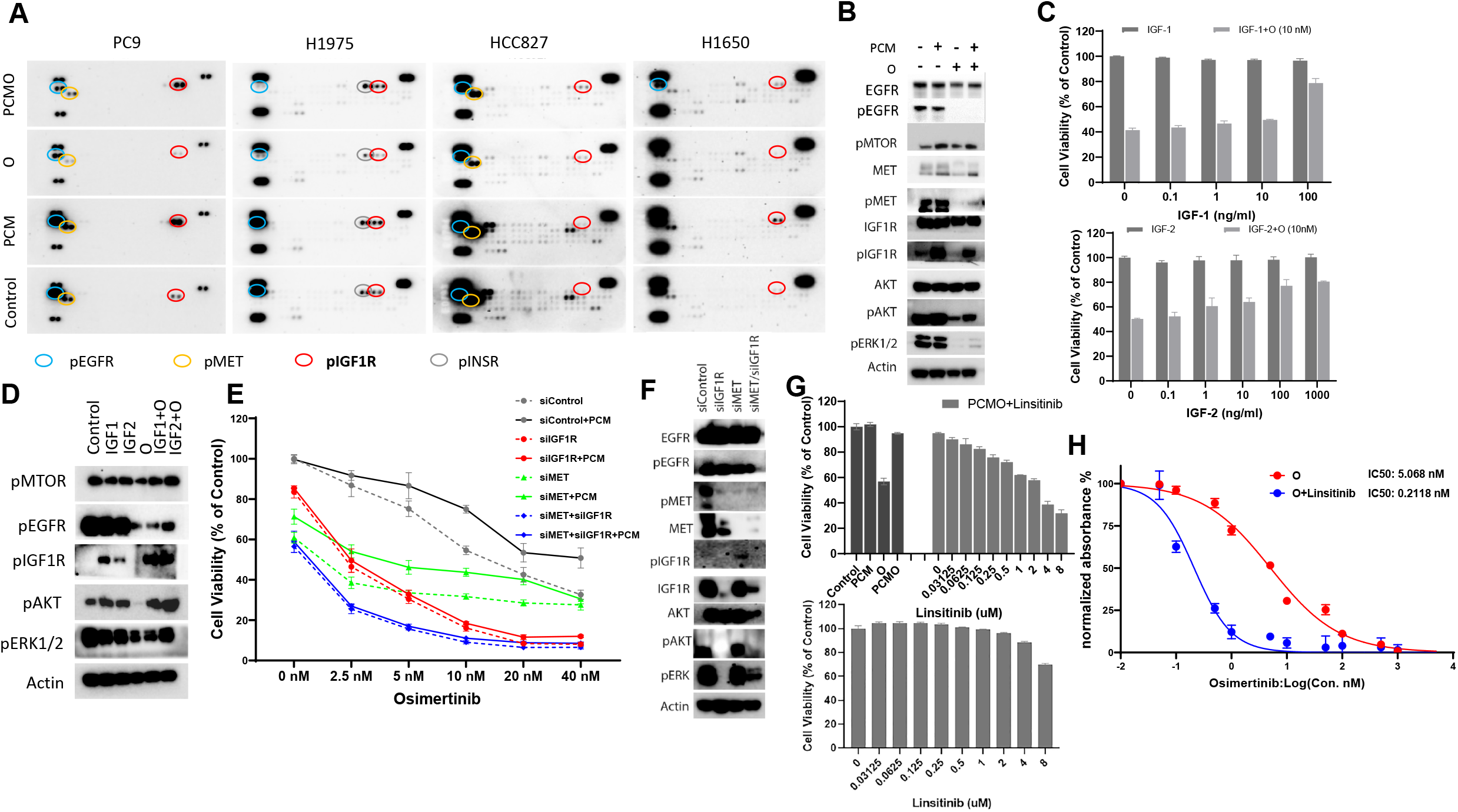
IGF1R bypass pathway activation contributes to osimertinib resistance in EGFR-mutant lung adenocarcinoma cells. (A) Phosphorylation of a panel of 49 RTKs was tested in a RTK array. PC9, HCC827, H1650 and H1975 cells, were treated with osimertinib (50 nM, 25 nM, 5 uM and 5 uM, respectively and *C. indologenes* PCM (100x dilution) for 24h. (B) Western blot of indicated proteins in PC9 cells treated with osimertinib (50 nM) and PCM (100x dilution) for 24h. (C) CCK-8 cell viability assay following treatment with indicated concentrations of IGF1 and IGF2 in PC9 cells. The assay was performed after 3 days of co-incubation. (D) Western blot of indicated proteins in PC9 cells after cells were treated with osimertinib (50 nM) and IGF1 and IGF2 (100 ng/ml) for 24h. (E) Cell viability assay of PC9 cells with knockdown of IGF1R (siIGF1R), MET (siMET) or both IGF1R and MET (siIGF1R +siMET). Cells were treated with osimertinib with or without *C. indologenes* PCM. (F) Western blot of indicated proteins in PC9 cells treated with siIGF1R, siMET for 24h. (G) CCK-8 cell viability assay following treatment with indicated concentration of osimertinib (10 nM) and linsitinib in PC9 cells, (H) Drug sensitivity assay was performed after 3 days treatment with osimertinib in siIGF1R-PC9 and siMET-PC9 cells.

Finally, we observed a bidirectional interaction between PC9 cells and *C. indologenes*: conditioned medium from PC9 cells significantly enhanced *C. indologenes* growth *in vitro* (p < 0.01) (Fig. 3E, F), while *C. indologenes* PCM increased PC9 cell viability and proliferation under osimertinib exposure. These results support the existence of a positive feedback loop between LAC cells and *C. indologenes* that could promote tumor persistence during targeted therapy. Future work should assess whether this growth-promoting effect is specific to *C. indologenes* compared to other bacterial species, thereby confirming the specificity of this cancer-microbiome interaction.

### Activation of the IGF1R signaling pathway by C. indologenes confers resistance to osimertinib in EGFR-mutant LAC

To investigate the mechanism by which *C. indologenes* mediates resistance to osimertinib in EGFR-mutant LAC cells, a phospho-RTK array was utilized to assess phosphorylation changes in 49 RTKs in PC9 cells treated with *C. indologenes* PCM. As depicted in Fig. 4A, PCM did not alter EGFR phosphorylation levels but significantly elevated the phosphorylation of IGF1R and c-Met, both in the presence and absence of osimertinib. These findings were corroborated by Western blot analysis (Fig. 4B), and enhanced phosphorylation of IGF1R and c-Met was similarly observed in H1975, H1650, and HCC827 cells (Fig. 4A).

To determine whether IGF1R pathway activation contributes to osimertinib resistance in EGFR-mutant LAC cells, PC9 cells were treated with IGF1 and IGF2 proteins. Both ligands increased cell viability relative to osimertinib monotherapy (Fig. 4C) and induced significant phosphorylation of IGF1R, mTOR, and ERK (Fig. 4D). Knockdown experiments using siRNA targeting IGF1R or cMet demonstrated that depletion of either receptor sensitized PC9 cells to osimertinib (Fig. 4E), with Western blots confirming efficient IGF1R knockdown (Fig. 4F). Interestingly, IGF1R silencing did not affect phosphorylated ERK (pERK) or AKT (pAKT) levels, implying PCM may stimulate alternative receptors or pathways; however, this residual signaling was insufficient to confer osimertinib resistance.

Further analysis revealed that IGF1R knockdown abolished PCM’s ability to rescue PC9 cells from osimertinib-induced cytotoxicity, whereas cMet depletion did not prevent PCM-mediated viability increases. IGF1R depletion reduced phosphorylation of AKT, mTOR, cMet, and nearly eliminated ERK phosphorylation; conversely, cMet depletion did not alter ERK phosphorylation (Fig. 4F). Co-treatment with the IGF1R inhibitor linsitinib and osimertinib, in the presence of PCM, significantly increased cell sensitivity to osimertinib and decreased PCM’s protective effect (Fig. 4G, H). These results collectively indicate that *C. indologenes* promotes osimertinib resistance primarily through activation of the IGF1R signaling pathway.

### Proteins secreted by C. indologenes play a role in mediating resistance to osimertinib in EGFR-mutant LAC cells

To investigate whether proteinaceous components within *C. indologenes* PCM are responsible for mediating resistance to osimertinib in LAC cells, we conducted a series of biochemical characterizations. PCM was first subjected to proteolytic digestion with Proteinase K, after which its ability to attenuate osimertinib–induced cytotoxicity was completely abolished (Fig. 5A), indicating that proteins are essential for its activity. To assess the potential involvement of low-molecular-weight peptides, PCM was fractionated by ultrafiltration, and the <10kDa flowthrough fraction was isolated and tested. This fraction failed to confer any protective effect (Fig. 5B), suggesting that the active component(s) are greater than 10kDa in size. Furthermore, heat inactivation of PCM at 95°C for 30 min, a condition sufficient to denature most proteins but not typically disruptive to small heat-stable peptides, also resulted in complete loss of activity (Fig. 5C). Collectively, these findings demonstrate that the bioactive factor(s) in *C. indologenes* PCM responsible for mediating resistance to osimertinib are heat-labile proteins larger than 10kDa, and effectively exclude the involvement of small peptides or other non-proteinaceous molecules.

**Figure 5.**
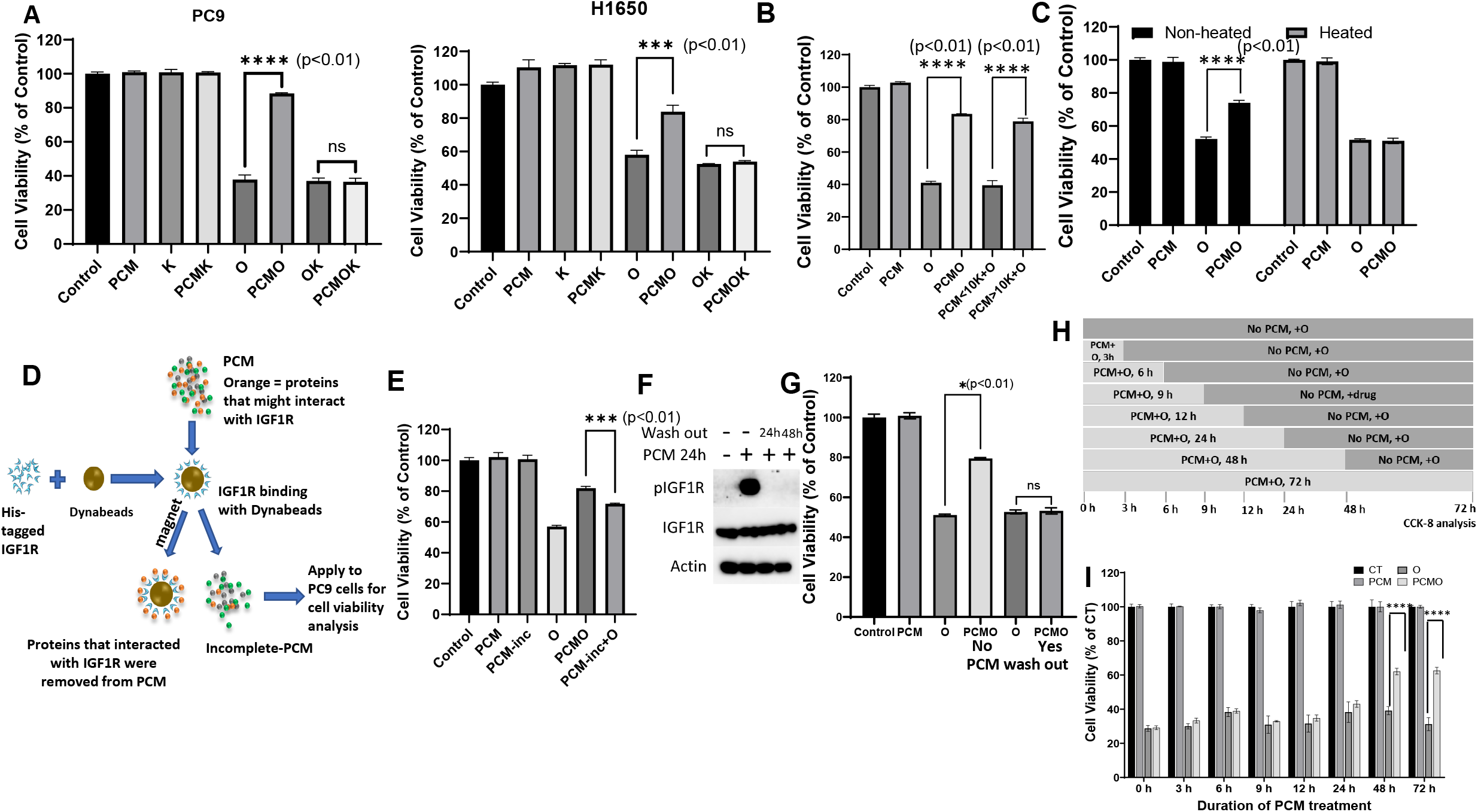
*C. indologenes* secreted proteins directly interact with IGF1R to activate IGF1R pathway. (A) CCK-8 assay following treatment with osimertinib, PCM and proteinase K in EGFR-mutant LAC cells. “O” = osimertinib, “K” = proteinase K. PCM was pre-treated with proteinase K for 30 min and then applied to LAC cells. (B) CCK-8 assay after treatment with PCM fractions (<10kDa or >10kDa) obtained by ultrafiltration. (C) CCK-8 assay following heat inactivation of PCM at 95°C for 30 min. (D) Schematic representation of the removal of proteins that might interact with IGF1R from PCM. After removal, the PCM was defined as PCM-incomplete. (E) CCK-8 assay following treatment with osimertinib, PCM and incomplete-PCM in PC9 cells. Here, PCM means that PCM was applied to beads exclusively without interaction with IGF1R protein. (F) Western blot of indicated proteins in PC9 cells after washing of PCM. (G) CCK-8 assay following treatment with osimetinib and PCM. PC9 cells were treated with PCM and osimertinib (10nM) for 24h, then PCM was washed out and the cells were cultured for an additional 48h with osimertinib (10nM). (H and I) The effect of duration of PCM treatment on susceptibility of PC9 cells to osimertinib. (H) The upper panel shows a schematic representation of the duration of treatment with PCM in PC9 cells, while (I) the lower panel shows cell viability after 72h of treatment.

Since our study demonstrated that activation of the IGF1R signaling pathway is a *C. indologenes*-mediated mechanism of resistance to osimertinib in EGFR-mutant LACs, we hypothesized that proteins secreted by *C. indologenes* directly interact with IGF1R and play a key role in activation of the IGF1R signaling pathway. To test this hypothesis, we first investigated whether removing proteins likely to interact with IGF1R from PCM would attenuate PCM-mediated resistance to osimertinib in EGFR-mutant LAC cells. We used His-tagged IGF1R protein bound to magnetic beads to remove proteins that might interact with IGF1R from PCM to create incomplete-PCM (Fig. 5D). PCM and incomplete-PCM were applied to PC9 cells treated with or without osimertinib. CCK-8 cell viability assay showed that cell viability with incomplete-PCM/osimertinib treatment was 70%, higher than with osimertinib alone (58%), but significantly lower than with PCM/osimertinib treatment (81%) (Fig. 5E). These results showed that incomplete-PCM could attenuate PCM-mediated resistance to osimertinib in EGFR-mutant LACs, meanwhile, indicating that proteins in PCM might directly interact with IGF1R protein. To better understand if proteins in PCM directly interact with IGF1R, we treated PC9 cells with PCM (no drug treatment) for 24h and then PCM was washed out and cultures maintained for another 24h and 48h. Our western blot assay also showed that PCM alone dramatically increased the level of phosphorylation of IGF1R after 24h of treatment, but this increased level completely disappeared when PCM was washed out for an additional 24h or 48h (Fig. 5F). Meanwhile, the CCK-8 cell viability assay also showed that PCM failed to rescue the PC9 cells from osimertinib when PCM was washed out after 24h of treatment (Fig. 5G). This indicated that IGF1R pathway activation occurs through binding to protein in PCM and not through the activation of endogenous IGF1R pathway activation factors. Based on these results, we believe that short-term treatment of PCM would not be sufficient to mediate resistance to osimertinib in LAC cells. To test whether the duration of treatment with PCM could affect the susceptibility of PC9 cells to osimertinib, we treated PC9 cells with the combination of PCM and osimertinib for 3, 6, 9, 12, 24, 48, and 72h, respectively (Fig. 5H), and then PCM was washed out, CCK-8 analysis was performed after 72h of treatment. As predicted by the removal experiment and western blot, treatment with PCM for 3 to 24h is not sufficient to affect cell susceptibility to osimertinib for up to 48h (Fig. 5I). Taken together, these results suggest that some specific proteins in PCM directly interact with IGF1R and activate IGF1R signaling pathway, thereby conferring resistance to osimertinib in EGFR-mutant LAC cells.

### *C. indologenes* increase resistance to osimertinib in fresh patient-derived EGFR-mutant LAC cells

To understand whether *C. indologenes* might also confer resistance to osimertinib in patient-derived EGFR-mutant lung adenocarcinoma cells, we derived two EGFR-mutant LACs, namely, COH4 (EGFR L858R) and COH561912 (EGFR L858R), from patients. To test the effect of PCM on cell viability of patient-derived LAC cells, we performed a CCK-8 assay in combination with osimertinib and PCM. Increased cell viability was observed when PCM was supplemented to osimertinib treatment in both COH4 and COH561912 (Fig. 6A). Half maximal inhibitory concentration (IC_50_) analysis showed that PCM significantly reduced cell sensitivity to osimertinib (Fig. 6B). This result indicated that PCM can induce resistance to osimertinib in patient-derived EGFR-mutant LAC cells. Since we identified activation of the IGF1R pathway as a mechanism mediating resistance to osimertinib in EGFR-mutant LAC cells, we assessed whether PCM also mediates resistance to osimertinib in patient-derived LAC cells through activation of the IGF1R pathway. RTK assay demonstrated that PCM can also increase IGF1R phosphorylation level in COH4, but not in COH561912 (Fig. 6B, C).

**Figure 6.**
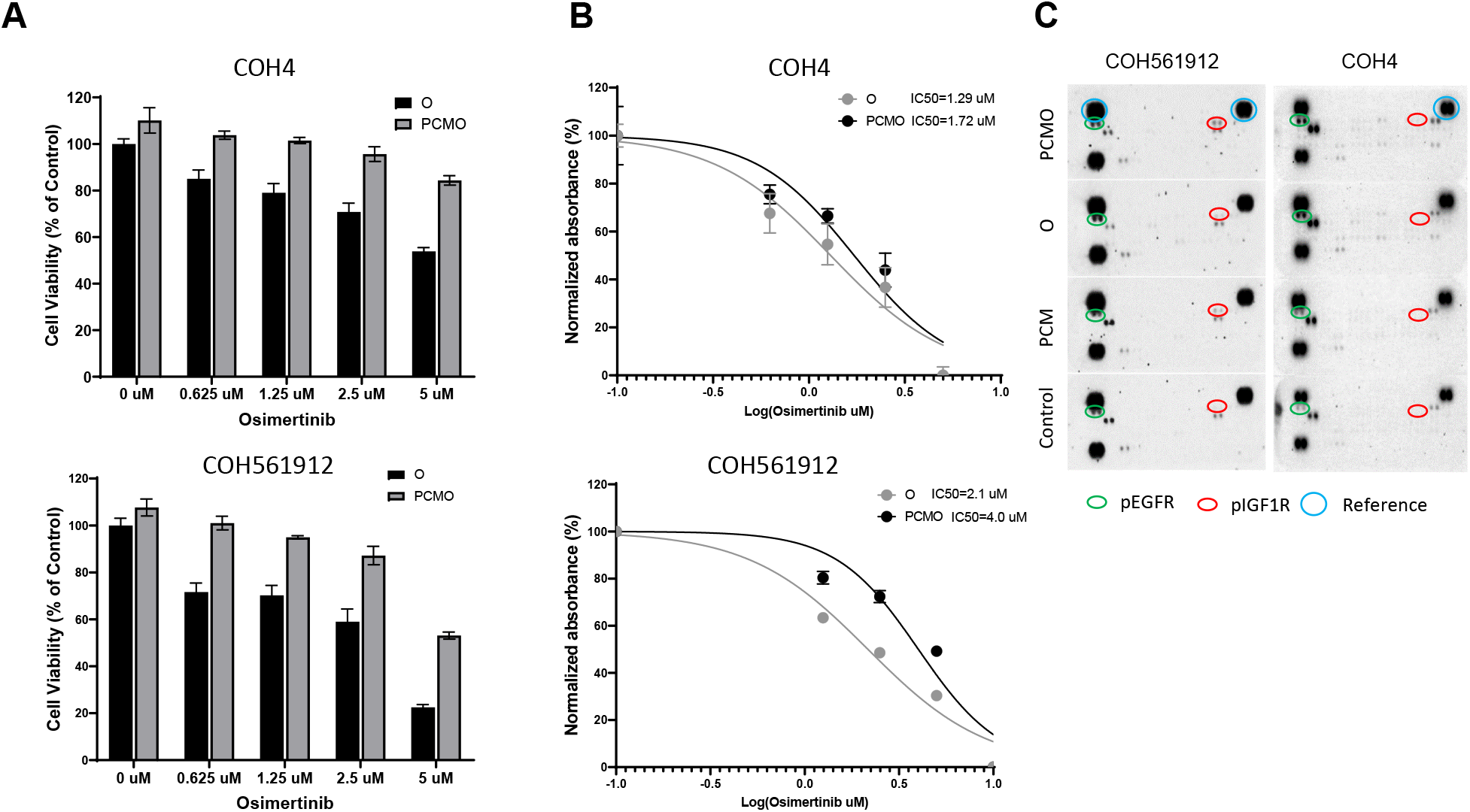
*C. indologenes* also mediate resistance to osimertinib in patient-derived EGFR-mutant lung cancer cells. (A) CCK-8 assays following treatment with indicated concentration of osimertinib and PCM in patient-derived EGFR-mutant lung cancer cells. (B) the IC_50_ of osimertinib in patients derived EGFR-mutant lung cancer cells. (C) Phosphorylation of RTK array in COH4 and COH561912, cells were treated with osimertinib (5µM) and PCM for 24h. (D) Western blot of indicated proteins in COH4 and COH561912 cells treated with osimertinib (5µM) and PCM (100x dilution) for 24h.

## Discussion

Although osimertinib is commonly used as first-line treatment for patients with advanced lung cancer with sensitizing EGFR mutations, patients eventually develop resistance. The mechanisms of acquired resistance to osimertinib are numerous, highly variable, and incompletely understood. Therefore, there is a desperate need for novel strategies to prevent or overcome therapy resistance in lung cancer. Intra-tumoral bacteria are increasingly proving to be the cause of therapy resistance in cancer.^32, 33^ In this study, we found that intra-tumoral *C. indologenes* confers resistance to EGFR-TKI therapy including osimertinib, and the presence of *C. indologenes* may decrease sensitivity to EGFR-TKIs. In addition, we investigated the mechanism by which *C. indologenes* mediates osimertinib resistance in EGFR-mutant LACs.

*C. indologenes* is normally found in soil, plants, and foods, but rarely in humans. Even though the clinical significance of *C. indologenes* has not been established due to infrequent recovery from patients, this species has been investigated in the context of squamous cell carcinoma of lung and nasal tube^38, 39^. *C. indologenes* has also been detected in human lung tumor specimens^32^, suggesting that humans could act as a host to this bacterium. In this study, we report, for the first time, that *C. indologenes* can decrease sensitivity and lead to resistance to osimertinib in EGFR-mutant LACs and fresh patient-derived lung cancer cells with EGFR mutation. In addition to *C. indologenes*, three of the other five species of the *Chryseobacterium* genus that we tested in this study also affected sensitivity to osimertinib in LACs to varying degrees. This implies that the presence of certain intra-tumoral bacteria, including *Chryseobacterium*, and potentially other related bacteria, can result in decreased sensitivity and even resistance to EGFR-TKIs in people with EGFR-mutant lung cancer.

Activation of the IGF1R signaling pathway has been reported to be a mechanism leading to acquired resistance to osimertinib in LACs.^5, 40^ However, in those studies, the IGF1R signaling pathway was activated by endogenous factors such as upregulation of IGF2 expression. Our findings also demonstrate that *C. indologenes* mediates resistance to osimertinib in LAC cells by activating the IGF1R signaling pathway. However, the activation of the IGF1R signaling pathway does not occur through endogenous factors, but rather through the direct interaction between IGF1R and proteins secreted by *C. indologenes* (Fig. 7). Our western blot and IGF1R knockdown results clearly demonstrate this mechanism. This proposed mechanism suggests the possibility of treating *C. indologenes*-mediated drug resistance by developing a treatment strategy that targets specific bacteria to improve susceptibility to osimertinib in EGFR-mutant LAC cells.

**Figure 7.**
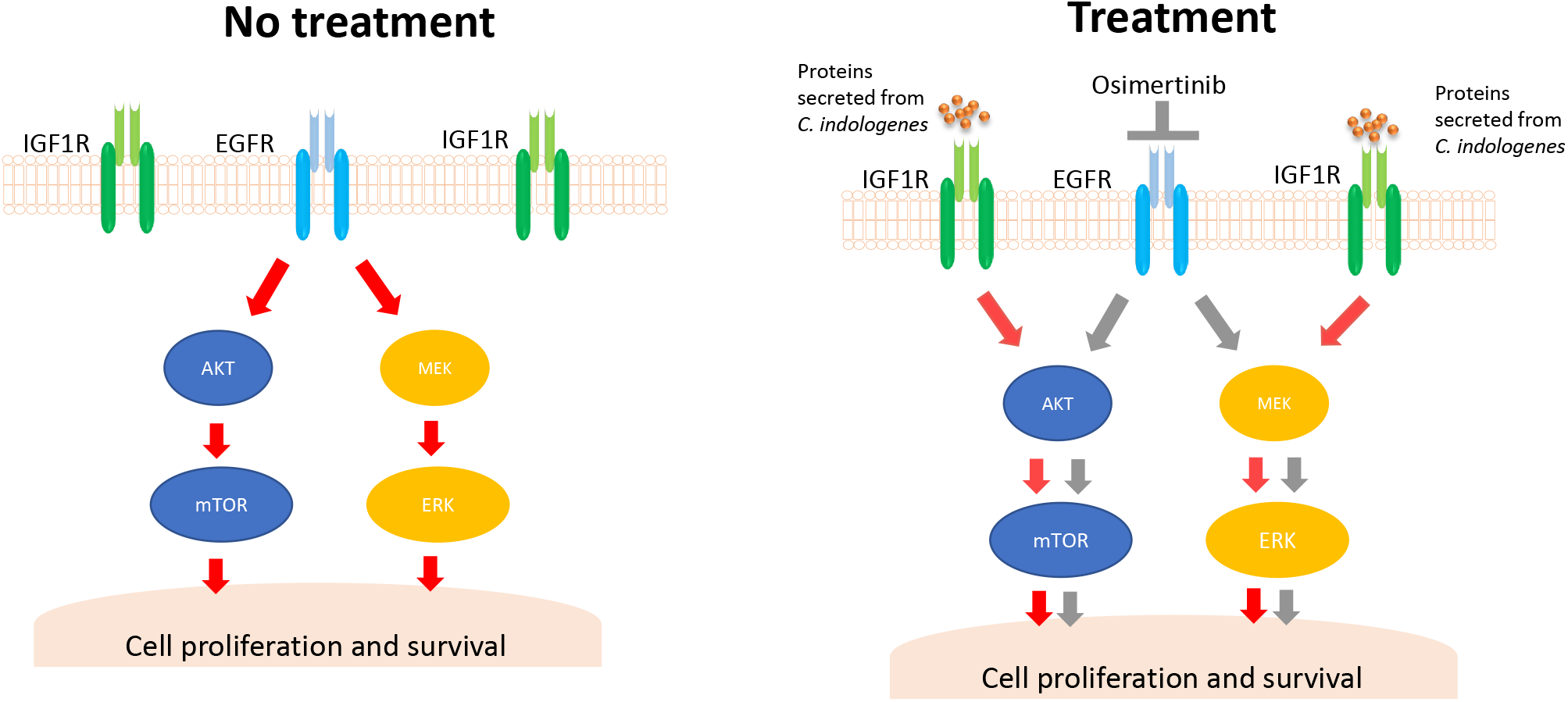
Schematic representation of the mechanism underlying *C. indologenes*-mediated osimertinib resistance in lung cancer cells. Proposed model of *C. indologenes* inducing an IGF1R bypass and contributing to acquired resistance to osimertinib in LAC cells. Red arrows depict activation of the pathway and grey arrows depict inactivation of the pathway.

Numerous studies have demonstrated the critical role that bacteria play in tumorigenesis^41^, antitumor immune response^42^, chemotherapy side effects^43^ and drug resistance^33, 44^. Our study presents a resistance mechanism by which *C. indologenes* directly activates the IGF1R signaling pathway in LAC cells. This differs from other reported mechanisms of bacteria-mediated chemotherapeutic resistance, such as metabolizing the chemotherapy drug gemcitabine (2,2-difluorodeoxycytidine) to its inactive form, 2,2-difluorodeoxyuridine. Our study also showed that LAC-conditioned medium can promote growth of *C. indologenes*, while *C. indologenes* could increase the viability of LACs, thereby suggesting a positive feedback loop between LAC cells and *C. indologenes*. This points to the potential role of the tumor-microbial microenvironment in anticancer drug resistance. Taken together, our results indicate that lung tumors harbor bacteria that can potentially impair the susceptibility of cancer cells to osimertinib, suggesting that intratumoral bacteria may contribute to anticancer drug resistance. Additional research is required to understand the prevalence of *C. indologenes* and related bacteria in lung cancers and how these may impact response to EGFR-TKI therapy in patients.

The results reported here could potentially lead to new interventions that may improve sensitivity to osimertinib or may lead to new therapies for patients with IGF1R-mediated osimertinib resistance. In addition, treatments that target specific bacteria may help better sensitize cancers to EGFR-TKIs and potentially improve the efficacy of first-generation generic EGFR-TKIs that are more affordable for communities with limited access to healthcare.

## Acknowledgements/Funding

Research reported in this publication was supported by National Cancer Institute of the National Institutes of Health under award number 1R21CA263751-1. Research reported in this publication also included work performed in core facilities supported by the NCI of the NIH under award number P30CA033572. The content is solely the responsibility of the authors and does not necessarily represent the official views of the National Institutes of Health. This work was also supported by the generous support of the Baum Family Foundation.

